# No Effect of Chronic or Acute Pain on Working Memory in the Sternberg Task

**DOI:** 10.1101/2025.05.13.653893

**Authors:** Caroline E. Phelps, Larissa E. Oliveira, Michelle N. Ngo, Kera S. Medhi, Stephanie Schoone, Devyn Debrosse, Lizbeth Aragundi, Alexe G. R. Delval, Robert C. Wilson

## Abstract

Memories of pain can last a lifetime, preventing future injuries, but this comes at the expense of remembering other concurrent experiences. For acute pain this cost is outweighed by the benefits but when pain is chronic, pain memory benefits are limited and may even contribute to maintenance of the disorder. Here we investigated two hypotheses, (1) pain takes up slots in working memory or (2) pain induces arousal above optimum levels at high task difficulty, leading to a decrease in performance. To do this, we used the Sternberg Task of working memory, in which the participant plays repeated trials where they are shown different sets of numbers and then asked to identify whether a ‘probe number’ was in the set or not. The Sternberg Task is ideal for testing the two hypotheses by looking at pain-related changes in accuracy and response time. There was a replication of the response time increase with set size, as well as the effect of older age on working memory and pain threshold. However, we saw no pain effect on either response time, accuracy, or the relationship between these parameters and set size with either chronic pain or an acute painful thermal stimulus. Together, this suggests that pain does not impair working memory in the Sternberg Task.

## Introduction

Memories of pain are vital for survival. When an individual touches a hot stove, reflexive withdrawal reduces injury in the moment, but associative pain memory protects against similar injuries for a lifetime. This retained memory may be crucial in older age, when avoiding unknown noxious stimuli is harder, due to an increase in pain thresholds[1, 2], decreasing the inherent warning signal of pain. However, strong pain memories come at the cost of remembering other concurrent experiences. Acutely the benefit of future protection from injury outweighs this cost. Yet, when pain is chronic, pain memories are detrimental, as inability to extinguish pain memories or fear of pain may lead to maintenance of chronic pain [3, 4] and the deficit in memories of other experiences can add to the debilitation of the disorder[5, 6]. Therefore, it is important to understand what underlies the memory deficits seen with chronic pain. Here, we particularly focus on working memory, which involves recalling and manipulating information over a short duration.

Deficits in working memory are frequently reported in chronic pain of many etiologies[7–9]. However, it’s complicated to determine whether pain itself or other aspects of the disorder in concert with/or excluding pain are to blame. Working memory deficits in chronic pain could result from changes in connectivity, function or structure of brain regions, such as loss of grey matter in the PFC[10, 11] (although see [12]), as well as medication use[13] or comorbidities like depression and anxiety[14, 15]. Therefore, to understand the role of pain in working memory deficits, there is a need to study pain alone, by applying acute pain to healthy volunteers.

Acute pain induced working memory deficits have been found in the n-back task only when the task load is high [16–18]. A prominent hypothesis for this load effect, is that pain uses up limited ‘slots’ in working memory[19, 20]. When task load is low, there are still sufficient ‘slots’ available for pain and task variables, but when task load is high, the concurrent pain leaves insufficient available for normal performance. However, using the n-back task alone cannot exclude other hypotheses for how pain causes working memory deficits. For instance, pain is inherently arousing and both too high and too low arousal induce non-optimal cognitive performance, seen by the inverted u-shape of the Yerkes-Dodson curve[21]. So, it could be hypothesized that when the task is difficult, pain shifts arousal to greater than optimal causing observed deficits. Most n-back task studies only use two working memory load conditions, perhaps due to task-induced fatigue[22] and improvement with practice[23], making it difficult to parse out a potential Yerkes-Dodson effect.

Therefore, to address these two hypotheses, we use the Sternberg task[24] under both acute and chronic pain conditions. In the Sternberg task the participant is shown a sequence of varying length (the set size), before identifying whether a ‘probe’ number was in the set. To test hypothesis 1, that pain takes up slots in working memory, this task is ideal because the longer the sequence (i.e. the more ‘slots’ used in working memory), the greater the response time. If pain is taking up slots in working memory, participants will be slower to respond on pain trials/conditions as more slots in working memory are used. Furthermore, this pain effect would be exacerbated in older individuals due to less initial cognitive resources. If hypothesis 2, that pain is arousing, is true then pain would induce a shift towards optimum arousal and improved performance variables at lower set sizes whilst pain would induce a shift away from optimum arousal and decrease in performance at higher set sizes.

## Methods

### Participants

#### Recruitment and Screening Procedure

##### Study 1

Study 1 tested the effect of applied thermal pain on both younger (18-30) and older (60-75) participants. Participants were recruited locally through a variety of methods including advertisements on the psychology department website, undergraduate listserv, posters around the university, at booths at local events, and via our internal call-back list from other in-lab projects. Participants were screened through an online REDCap[25] questionnaire and invited to the lab if eligible. Exclusion criteria included known neurological or cognitive disorders which could affect memory, taking centrally acting pharmacological agents which could affect pain thresholds (e.g. prescription analgesics and antidepressants) and chronic pain, metabolic or circulatory disorders. All participants gave informed consent and were paid $15 per hour, which was not contingent on performance.

##### Study 2

Study 2 tested young participants with chronic pain and healthy controls, without any external painful stimuli. Participants were undergraduate students at the University of Arizona who signed up for the study on the Psychology Department’s SONA system. Some participants were invited to sign up for the study based on their answers on the Graded Chronic Pain Survey-revised (GCPS-r) [26] completed in a course-specific mass survey, although generally all people with access to the SONA group were able to sign up. When they came to the lab they were screened using an online RedCap questionnaire, and exclusion criteria were significant neurological, psychiatric, medical illness or injury that would affect cognitive function. All participants gave informed consent and received course credit, which was not contingent on performance. Presence of chronic pain for all participants was determined by the GCPS-r[26], completed on the day of testing, in a Qualtrics questionnaire or written format.

All experiments were conducted at the University of Arizona, Tucson, USA and these studies were approved by the University of Arizona Institutional Review Board. This study represented preliminary data and was not pre-registered. No patients were consulted in the design of this research.

#### Sample Demographics

##### Study 1

Older participants had significantly higher education attainment than younger participants (Table 1, Kolmogorov–Smirnov test, D=0.65, p<0.001). A Chi-square test of independence found no significant differences in gender composition between age groups. However, there were significantly higher number of females in the younger participant group than expected proportions (χ^2^ = 4.67, p = 0.031).

**Table 1:**
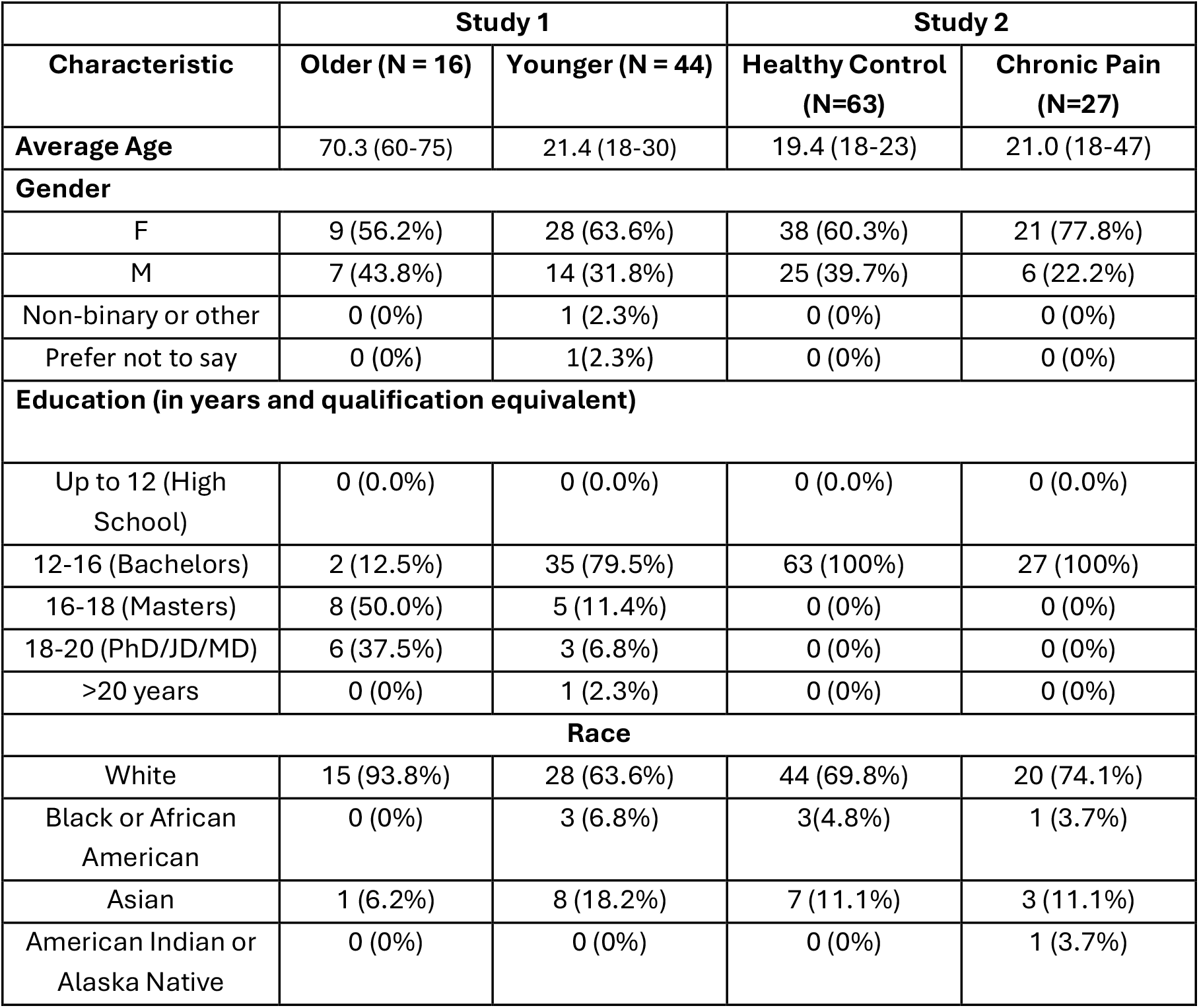
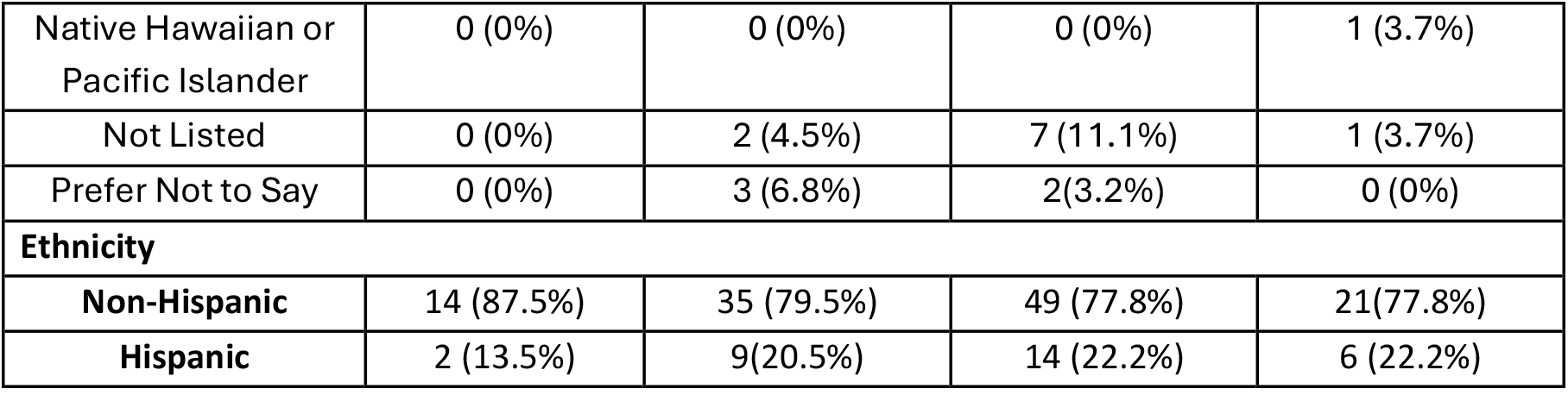
Demographics for all included participants.

16 (26%) younger and 11 (41%) older participants were excluded for having thresholds higher than maximum allowed temperature on both arms at time of testing. The maximum temperature was initially 48°C (n=27 younger) but raised to 50°C (n=27 older, n=33 younger) due to greater than expected younger participants exceeding the 48°C limit (7 (22%) exclusion, only younger tested). 1 younger participant withdrew after pain threshold testing due to pain-related anxiety.

##### Study 2

A Chi-square test of independence found no significant differences in gender composition between control and chronic pain groups and there was also no difference in education attainment as all subjects were in undergraduate programs (Table 2). 3 (4.5%) control participants were excluded from analysis for achieving less than 70% accuracy overall in the Sternberg working memory task, whilst no (0%) chronic pain participants were excluded.

**Table 2:**
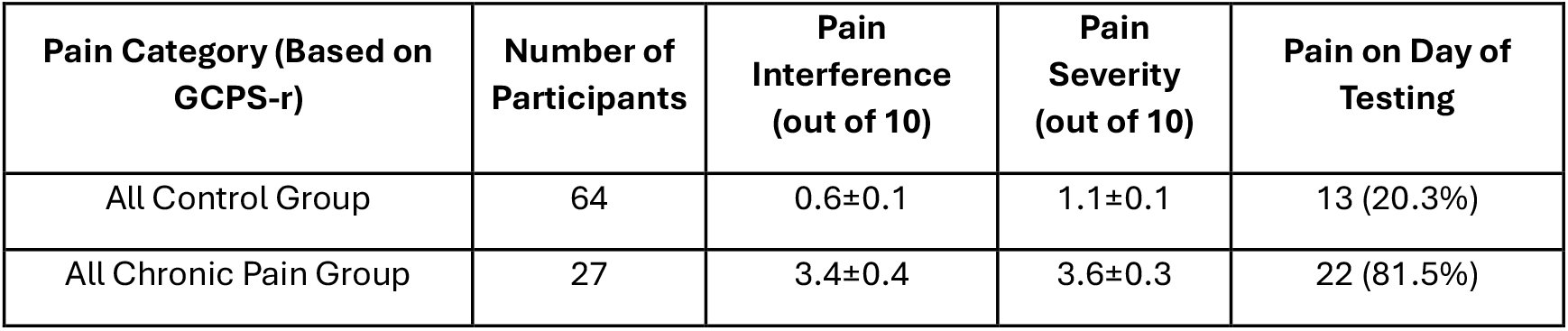
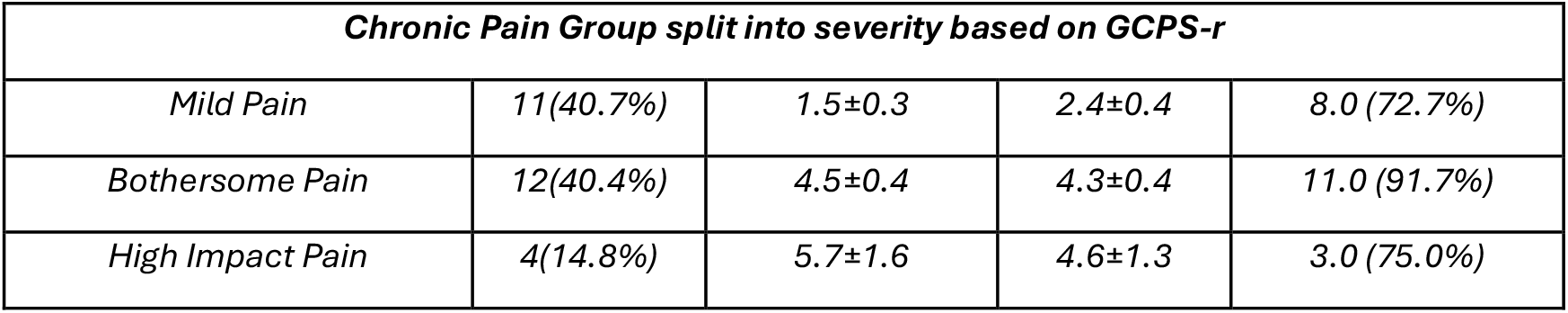
Severity, Interference and Pain in the Chronic Pain Group in Study 2. Severity of chronic pain classification is from the Graded Chronic Pain Scale-revised, and average pain severity and intensity as well as presence of pain on day of testing was from the Brief Pain Inventory – Short Form.

### Pain threshold testing (Study 1 Only)

Pain threshold testing and the Sternberg Working Memory task were coded and run in Psychopy (Study 1: version 2023.1.1, Study 2: version 2022.2.5 or 2023.1.1) [27]. Pain thresholds were determined using the adaptive staircase procedure [28]. After reading the instructions (based on [29]), participants placed their left volar forearm on the thermosensory stimulator (QST.lab, TCS II). Each stimulus was 8s, including 3s ramping between baseline and target temperature. After the offset of the heat stimulus, the participants rated their pain level on a sliding numeric rating scale (NRS) between 0 and 10, presented on the computer screen, where 0 was no sensation, 2 was considered pain threshold and 8 was pain tolerance. Four different areas on the left volar forearm were tested, with three rounds of heat at each site. The first three temperatures were the same for all participants (41°C, 44°C, and 47°C), then subsequent stimulus temperatures at the other 3 sites were selected using an iterative linear regression to identify temperatures predicted to elicit ratings of 2 (low pain) and 8 (high pain) for each subject. If participants 2 or 8 value exceeded the maximum allowed temperature, they were given the opportunity to repeat pain threshold testing on their right arm. Participants were excluded if their 2 or 8 temperature value was higher than the maximum temperature or their r^2^ value was lower than 0.4, on both arms.

### Sternberg Working Memory Task

On each trial of this task, participants observed a set of between two and six numbers. Each number appeared on the screen individually for 1.2s. After a delay of 2s during which a fixation cross is shown, they are then presented with a probe number and asked to respond as quickly as possible as to whether this probe was in the set of remembered items or not. There was a time-out of 3s.

#### Study 1

There were three randomized pain conditions: no pain, low pain (2 on NRS), high pain (8 on NRS). Pain was applied during the delay and response period (5s). To ensure equal heat application time, if the participant responded before 3s in the response period, there was a wait message and feedback would be presented after the 3s time period was finished. There were 4 blocks in the task and the location of stimulation on the forearm was moved between blocks to prevent sensitization or adaption to the thermal stimulus.

#### Study 2

No painful stimuli were applied. Participants had completed a one-hour decision battery prior to starting the Sternberg working memory task. Participants were excluded if they achieved less than 70% overall accuracy.

### Statistics

For all analyses, a linear mixed-effect model was fit using the lmer function in R (version 4.4.1). This was followed by a Type III ANOVA to assess significance of main effects and interactions and where significant effects were found, post-hoc pairwise comparisons of estimated marginal means were conducted, with a Tukey test for multiple comparisons.

For pain thresholds in study 1 the dependent variable was the temperature, and independent variables were pain rating, age group and subject. For Sternberg analyses, the dependent variable was response time or accuracy, and the independent variables were group (age or chronic pain status), set size (number of items in the sequence), subject and in study 1, the pain level applied (0 2 or 8). All figures were created in Prism (version 8).

## Results

### Study 1

#### Older Adults within Allowed Parameters Had a Higher Pain Threshold than Younger Adults

Initially pain threshold (rating of 2) and tolerance (rating of 8) were calculated using an adaptive staircase procedure on the participant’s forearm. Including all participants tested, apart from those who were excluded when maximum temperature was 48°C (Figure 1A), there was no significant effect of Age or Age*Pain Rating, suggesting that age doesn’t affect either the pain threshold or tolerance, or the relationship between the two. However, this may be a limitation of the experimental procedure. In determining pain thresholds, temperatures over 50°C (or 48°C, 27 younger participants) could not be applied, which reduced the ability to adaptively calculate thresholds over this temperature. Participants excluded when 48°C was the maximum, had particularly unrealistic computed tolerance, which is why they are excluded from the figure and analysis. Therefore, overall, pain tolerance may be particularly underestimated, limiting conclusions that can be drawn about age and pain tolerance. Especially considering that the percentage of participants who had a pain threshold or tolerance over allowed temperatures was 26% of younger participants but 41% of older participants.

**Figure 1:**
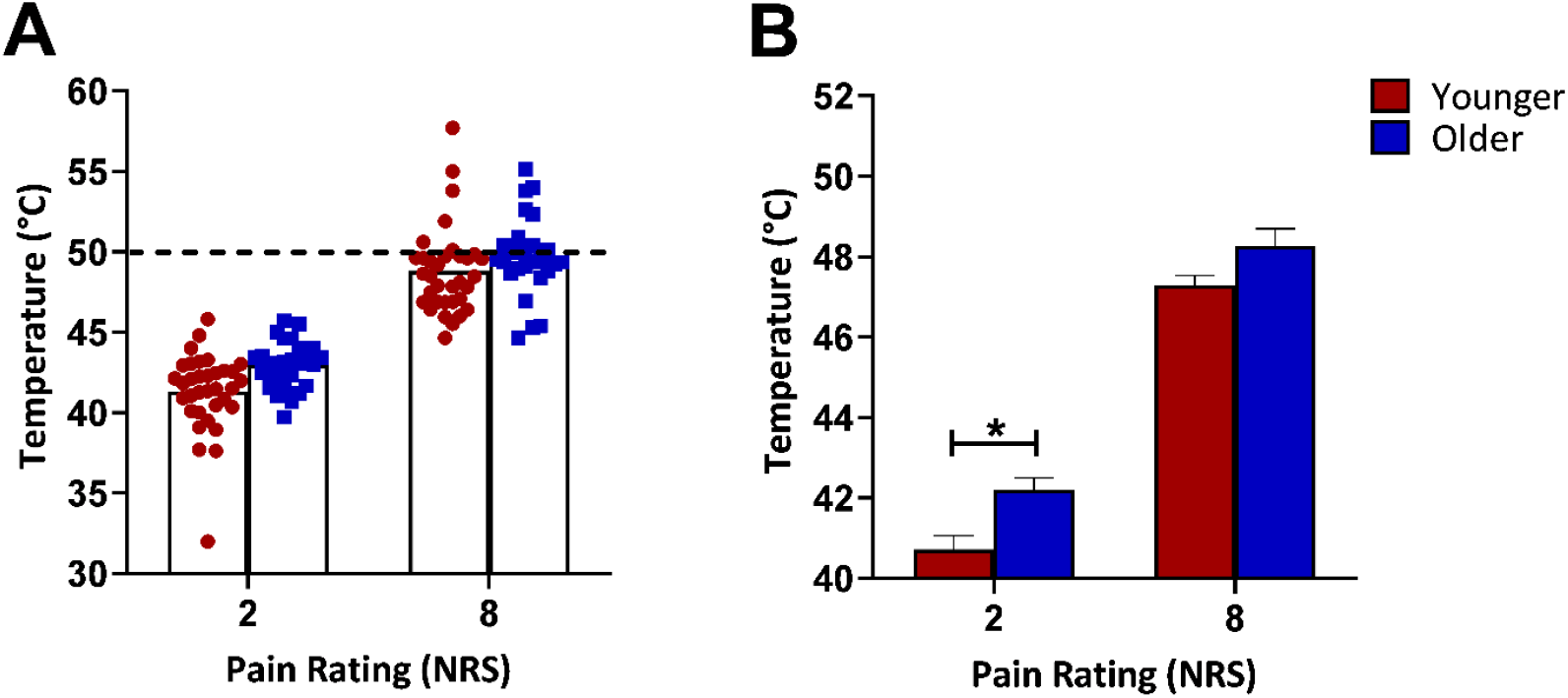
Estimated temperature eliciting a response of 2 or 8, in younger and older participants. (A) Individual pain thresholds (NRS = 2) and pain tolerance (NRS = 8) before exclusions. (B) Average pain thresholds and pain tolerance for participants after excluding participants whose pain tolerance was above 50°C. Participants excluded when 48°C was the maximum allowed temperature are not included in Figure 1A. * p<0.05 NRS: numeric rating scale

Estimated temperatures of pain threshold and tolerance of all participants who were under the maximum allowed temperature are shown in Figure 1B. These participants’ data are used in the Sternberg analysis in Figure 2. Interestingly, in comparison to the whole sample, in these included participants there was a significant effect of age (F(1,107.89) = 7.40, p=0.0075) with a higher pain threshold in older participants (t(108) = 2.72, p = 0.037) but no significant difference in pain tolerance.

**Figure 2:**
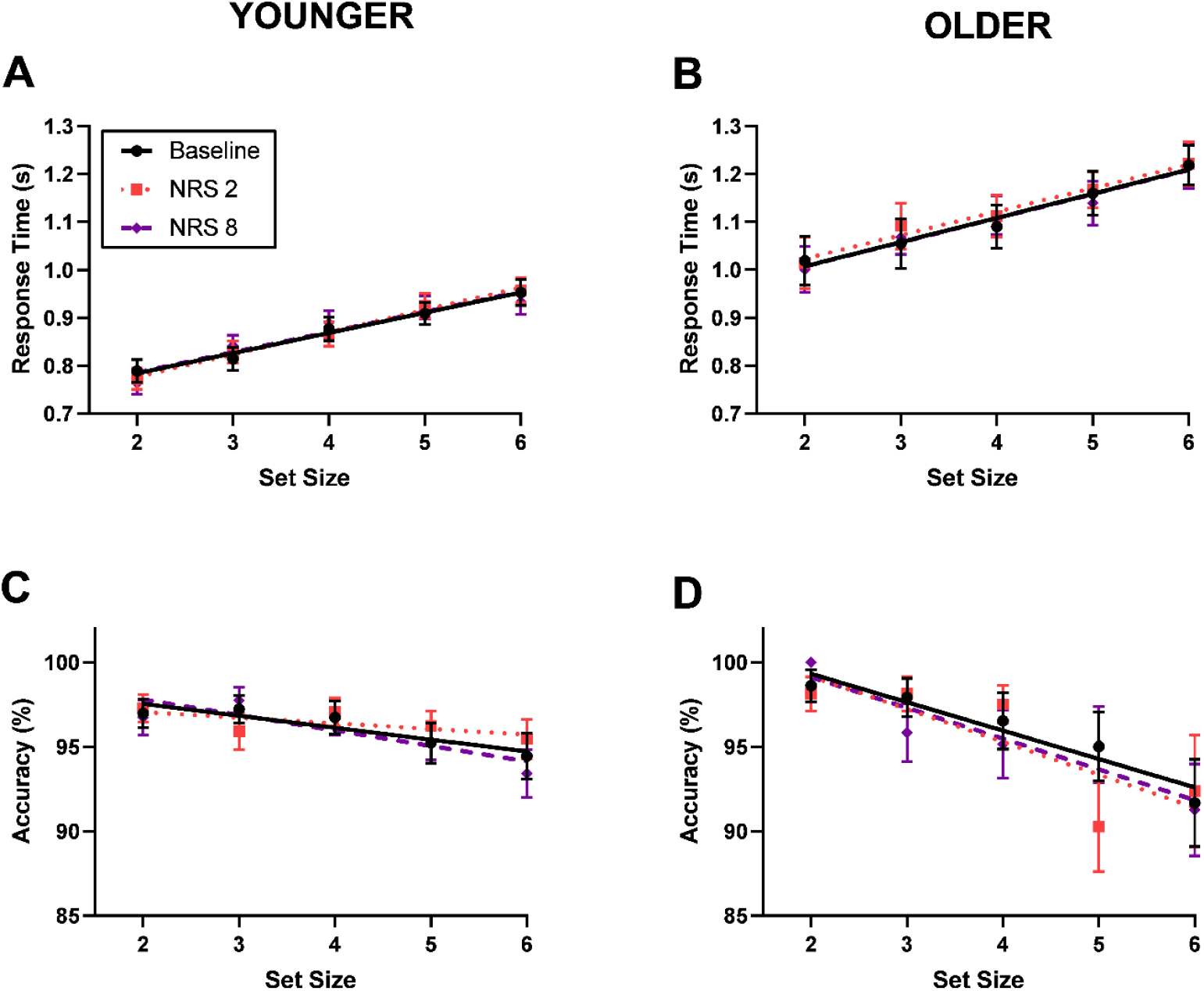
Response Time and Accuracy in relation to set size (number of items in the sequence) in participants under baseline (non-painful heat), NRS 2 and NRS 8 (the temperatures expected to elicit a rating of 2 or 8 for the participant on the numerical rating scale). Response time increases as a function of set size but there is no effect of pain in (A) younger or (B) older participants. There is also no effect of pain on accuracy in (C) younger or (D) older participants but older participants have a steeper decrease in accuracy with increasing set size.

#### Older participants have an overall slower response time and a larger decrease in accuracy as set size increases, but there are no effects of acute thermal pain

In the Sternberg Task of working memory, overall, there was a replication of the classic ‘Sternberg Effect’, in which an increase in set size leads to a longer response time. Younger participants here had a similar increase in time per item, 42.2 ± 7.7ms compared to the original young participants in Sternberg’s experiment who increased response time by 37.9 ± 3.8ms per item. Looking at both older and younger participants, the linear relationship between set size and response time was significant, with a main effect of set size (F(4, 812) = 175.56, p<0.0001, Figure 2A and B).

Older participants were slower overall, with a significant main effect of age (F(1,58) = 29.79, p<0.0001) and a significantly slower response time at every set size (Figure 2B). However, there was no change in the relationship between set size and response time, seen by no age*set size interaction and a non-significant difference in slope, illustrative of no change in time per item (baseline temperatures, older 50.5 ±14.7, younger 42.2 ± 7.7ms). This suggests a slower baseline reaction time, but not a bigger increase in response time with a greater set size. Despite similar findings to Sternberg[24] and an age-related effect on response time, suggesting a task concordant with the literature as well as amenable to individual differences, we see no effects of acute thermal pain. Neither pain applied at threshold (NRS 2) nor at higher, tolerance levels (NRS 8) had any effect on response time or the relationship between response time and set size, in either younger or older participants.

The initial Sternberg paper did not find significant changes in accuracy with set size. However, here overall there was a decrease in accuracy with an increase in set size (F(4,812) = 11.52, p<0.0001, Figure 2C and D). This effect was more pronounced in older age, with a greater decrease in accuracy, the larger the set size (age*set size F(4, 812) = 3.09, p=0.015, Figure 2D) and a trend to significantly lower accuracy in older adults at the highest set size (t(168) =−1.846, p = 0.066). Similarly to response time results, neither pain level had any effect on either accuracy or the relationship between accuracy and set size.

### Study 2

#### Participants classed as having chronic pain from the Graded Chronic Pain Scale-revised (GCPS-r) reported significantly higher incidence of pain on day of testing, pain interference and pain severity

To determine if participants had chronic pain, they completed the GCPS-r which allowed us to categorize them into control and chronic pain groups. They also completed the Brief Pain Inventory short form (BPI-sf) for further information on pain severity and interference in their everyday life. There was a significant difference between controls and chronic pain for pain severity (t(88) = −8.16, p<0.001) and pain interference (t(88) = −8.04, p<0.001) as determined using the BPI-sf (Table 2). Participants with chronic pain also reported pain on the day of testing significantly more than controls (χ^2^ = 24.55, p<0.001, Table 2). Table 2 also shows the number of participants in each severity category, as defined by the GCPS-r. Over 80% of participants in the chronic pain group had either mild or bothersome chronic pain.

#### Participants with Chronic Pain did not differ in performance measures in the Sternberg Task of Working Memory

Similar to study 1, there was a replication of the classic Sternberg effect in which response time increases in proportion to set size (set size (F(4, 352) =138.14, p<0.0001, Figure 3A)). Here the time for each item in working memory was 51.2ms ± 2.5 for controls and 44.5ms ± 4.0 for chronic pain, slightly higher than the 37.9ms ± 3.8 reported in Sternberg’s original paper, but similar to times seen in study 1. However, there was no difference in the Sternberg effect in participants with chronic pain. This lack of change in either response time or relationship of response time and set size, indicates that there was no influence of chronic pain on either baseline reaction time or slots taken up in working memory.

**Figure 3:**
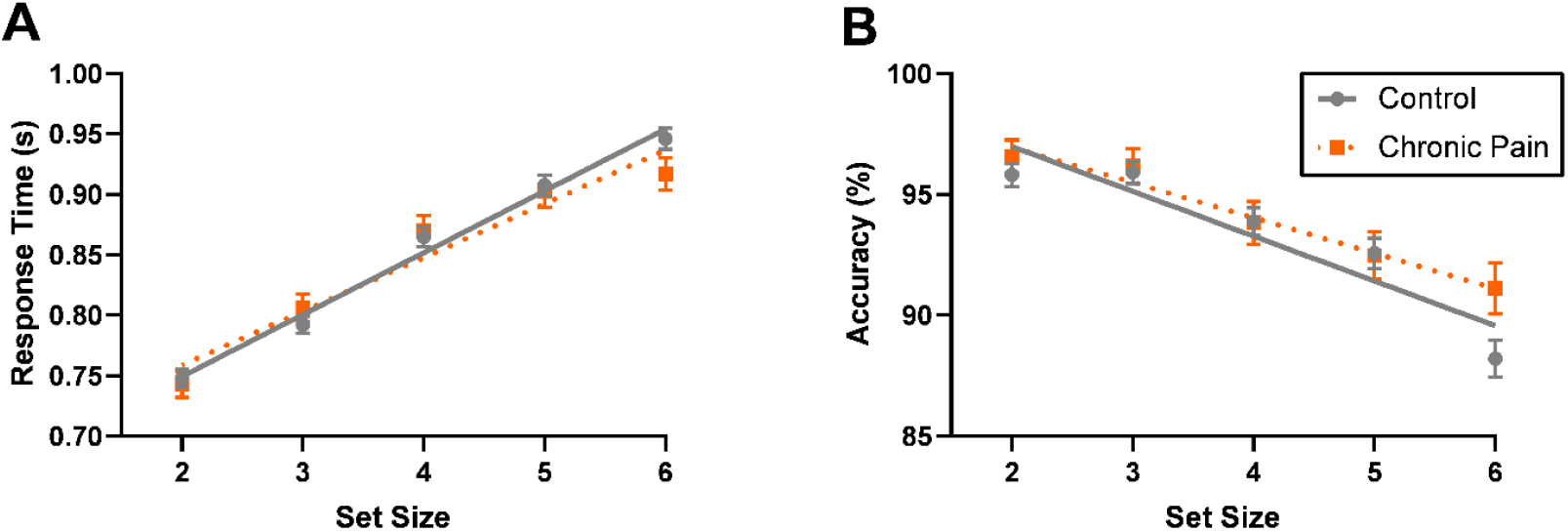
Neither response time (A) nor accuracy (B) differ between control and chronic pain groups. Response time increases as a function of set size and accuracy decreases as a function of set size in both groups.

Accuracy significantly decreased with set size (F(4,352) = 29.30, p<0.0001, Figure 3B), replicating the effect on accuracy seen in study 1. Chronic pain had no effect on accuracy or the set size accuracy interaction. This suggests that whilst accuracy reliably decreases with a greater number of items in working memory, chronic pain has no influence on this relationship.

## Discussion

Pain is purported to cause deficits in working memory, but the mechanism underlying this is unknown. We used the Sternberg task to differentiate between two hypotheses 1) pain induces deficits by taking up limited ‘slots’ in working memory, leaving insufficient available for the task and 2) pain causes arousal, shifting participants over optimum arousal states for performance. However, surprisingly, we saw no deficits in working memory either with acute pain or in participants with chronic pain, raising questions about the relationship between pain and working memory.

A possible explanation for negative results can be poor experiment design, which is unable to capture treatment or group differences. However, here we argue this is not the case, because we replicate several findings reported in the literature including the Sternberg effect, thermal pain thresholds, as well as the effect of older age on both. First, we replicated the ‘Sternberg effect,’ an increased response time with greater number of items in working memory. In the Sternberg task, participants see a set of numbers of varying length, then identify whether a ‘probe’ number was in the set. The increased response time with greater set size is thought to represent the effects of higher working memory load [30, 31]. Here, in both studies, the increase in time for each additional item in working memory was similar to Sternberg’s original paper (37.9ms per item)[24], suggesting a robust replication of his original findings. The only difference is that in Sternberg’s original paper no effect of set size on accuracy was reported[24], yet here accuracy decreased with increased set size in both studies.

Secondly, we replicated age group differences in both response time and accuracy. Older adults had significantly slower response time than younger adults, driven by an increase in the intercept, but not the slope, of the set size by response time plot. This suggests that older adults have a slower baseline reaction time compared to younger counterparts, rather than altered strategy or time to process each item in working memory. In contrast, accuracy had a steeper decline in older adults compared to younger adults with increasing set size. Thus, whilst older adults use similar working memory strategies and have a similar processing time to younger adults, they are more likely to make errors as the working memory load gets higher, potentially suggesting reduced working memory capacity or fidelity. This is in accordance with age-related deficits in working memory seen in other studies[32–34] and illustrates that the Sternberg task is sensitive to common age-group differences.

With regards to replicating classic pain effects, we saw increased thermal pain threshold in older adults. Whilst there was no significant increase in pain tolerance, this was likely because a greater number of older than younger adults were excluded from the study for having pain tolerance above temperatures allowed to be applied. In younger adults, similar threshold and tolerance was seen here as in 342 participants in [28], suggesting a good replication of this technique which has been used in other working memory studies (e.g. [16]). The higher pain threshold in older adults is also in accordance with a systematic review and is within the expected range for the age difference[1]. Although pain tolerance conclusions were again limited by the maximum allowed temperature. Overall, the pain threshold and tolerance results are in accordance with the literature, suggesting that we elicited pain during the Sternberg task at temperatures consistent with other publications.

Together, these replications of the Sternberg effect, pain and aging results suggest an experiment consistent with the literature and sensitive to age-group differences. Despite this, we saw no impact of pain on working memory in either trials with concurrent acute thermal pain or in a separate study comparing chronic pain to healthy controls. What does this mean for our initial mechanistic hypotheses? If the first hypothesis were correct and pain caused deficits by taking up slots in working memory, there would have been an increase in response time at each set size, and proportional to the intensity of pain, with higher intensity pain taking up more slots and causing a slower response time. In terms of accuracy, there would have been no effect on accuracy at smaller set sizes, where there is sufficient capacity for both pain and task in working memory, but a decline in accuracy at larger set sizes when working memory capacity is insufficient for both task and pain. If the second hypothesis were correct and pain was arousing, there would have been a faster response time and better accuracy than control at smaller set sizes and slower response times and decreased accuracy at larger set sizes, like the inverted u-shape of the Yerkes Dodson curve. However, instead there was no difference in response time or accuracy at any set size, neither in younger nor older participants when low or high intensity acute pain is compared to baseline heat trials, nor when participants with chronic pain are compared to healthy controls. There was even no effect of acute pain in older adults who had a steeper decline in accuracy with larger set sizes, which may be indicative of a reduced working memory capacity. This suggests that neither acute nor chronic pain have any effect on working memory performance in the Sternberg Task.

On the face of it, no effect of acute or chronic pain on working memory seems to be in contrast with prevalent ideas of pain induced cognitive deficits. However, this finding adds to the growing literature that suggests the relationship between pain and working memory is not clear cut. Where acute pain-induced deficits in working memory have been found, they are often in the n-back task[16, 17, 35]. Conversely, working memory deficits have not been found in tasks akin to the Sternberg task [17, 36], or only found when inter-individual factors such as pain catastrophizing are accounted for [37]. Yet even within the n-back task there are differences, the load required for pain-induced deficits is inconsistent(i.e. 2 or 3 back[16, 17, 35]) and conversely, higher load could be protective against pain seizing cognitive resources[38]. With regards to chronic pain, two meta-analyses have reported a moderate effect on working memory[7, 9]. However, many of the included chronic pain conditions are nociplastic, such as fibromyalgia in which there is a higher incidence of comorbidity with depression and anxiety than most other pain disorders [39], so this raises questions over the specificity of the deficit in working memory to pain. Therefore, the lack of effect of acute or chronic pain on Sternberg task performance seen here is a valuable addition to the growing literature, suggesting that pain alone is insufficient to alter working memory performance, except in particular types of tasks or with comorbidities.

Where the pain and working memory literature is consistent, is in load-dependent working memory induced analgesia. Analgesia occurs only at a high acute pain intensity and when the task is difficult [22–24], this is in line with hypothesis 1, suggesting that the trade-off between task and pain only occurs when there are insufficient cognitive resources for both. This task induced analgesia could lead to decreased interference of the painful stimulus on working memory because deficits in working memory depend on the amount of pain perceived in that moment, rather than how the pain was rated outside of task conditions [40]. Therefore, this may be one of the inherent difficulties of studying pain in the laboratory. Outside of laboratory conditions, pain indicates injury that needs to be escaped or recovered from[41], then remembered for avoidance in the future[42]. In contrast, in the lab, participants are informed that the heat applied cannot cause injury, so reducing inherent motivational drive prioritizing pain over task stimuli[41]. Therefore, acute pain studies in the laboratory may be of limited assistance to understanding cognitive deficits in chronic pain.

We saw no deficit in Sternberg task performance in chronic pain participants, in line with chronic low back pain [43]. However, a limitation is that our chronic pain group has heterogenous pain and their average pain interference and severity scores are relatively low, in comparison to those typical in a pain clinic environment. Yet, 81.5% of them experienced pain on the day of testing, suggesting they are a valid group for determining the impact of ongoing pain on working memory. However, maybe this group does not have sufficiently intense pain to see cognitive deficits, similar to the finding that chronic pain patients with milder pain did not show deficits in a difficult attention task, whilst those with higher intensity pain did[44]. In future work, comparing performance between high impact and mild chronic pain is needed, unfortunately we were unable to do that here because of the small sample size.

In conclusion, whilst pain is purported to cause deficits in working memory, we found no evidence of this in either acute or chronic pain in the Sternberg Task. These results add to the increasing number of reports suggesting that the relationship between pain and working memory may be more complicated than generally assumed.

## Acknowledgments

This study was partially funded by preliminary study funds from the pilot award P30 NIH-NIDA (3042525) Core Center of Excellence in Addiction Studies.

## Notes

### Competing Interest Statement

The authors have declared no competing interest.

